# Multi-centre, multi-vendor 7 Tesla fMRI reproducibility of hand digit representation in the human somatosensory cortex

**DOI:** 10.1101/2021.03.25.437006

**Authors:** Ian D Driver, Rosa M Sanchez Panchuelo, Olivier Mougin, Michael Asghar, James Kolasinski, William T Clarke, Catarina Rua, Andrew T Morgan, Adrian Carpenter, Keith Muir, David Porter, Christopher T Rodgers, Stuart Clare, Richard G Wise, Richard Bowtell, Susan T Francis

## Abstract

Whilst considerable progress has been made in using ultra-high field fMRI to study brain function at fine spatial resolution, methods are generally optimized at a single site and do not translate to studies where multiple sites are required for sufficient subject recruitment. With a recent increase in installations of human 7 T systems, there is now the opportunity to establish a framework for multi-site 7 T fMRI studies. However, an understanding of the inter-site variability of fMRI measurements is required for datasets to be combined across sites. To address this, we employ a hand digit localization task and compare across-site and within-site reproducibility of 7 T fMRI to a hand digit localization task which requires fine spatial resolution to resolve individual digit representations. As part of the UK7T Network “Travelling Heads” study, 10 participants repeated the same hand digit localization task at five sites with whole-body 7T MRI systems to provide a measure of inter-site variability. A subset of the participants (2 per site) performed repeated sessions at each site for measurement of intra-site reproducibility. Dice’s overlap coefficient was used to assess reproducibility, with hand region inter-site Dice = 0.70±0.04 significantly lower than intrasite Dice = 0.76±0.06, with similar trends for the individual digit maps. Although slightly lower than intra-site reproducibility, the inter-site reproducibility results are consistent with previous single site reproducibility measurements, providing evidence that multi-site 7 T fMRI studies are feasible. These results can be used to inform sample size calculations for future multi-site somatomotor mapping studies.

## Introduction

Functional magnetic resonance imaging (fMRI) provides a unique, non-invasive tool for resolving spatial features of neuronal modulations in the human brain, this is widely used to test hypotheses across the whole field of neuroscience research (Logothetis, 2008; Rosen and Savoy, 2012). The fMRI variant that shows the highest sensitivity without use of an exogenous contrast agent is the blood oxygenation level dependant (BOLD) contrast (Ogawa *et al.*, 1990, 1992; Kwong *et al.*, 1992).

BOLD fMRI can be performed at ultra-high field, higher field strengths than used for the majority of clinical workload (1.5-3 T), with the main advantage being finer spatial resolution. Increased sensitivity is available at high field, which can be traded for smaller voxel volumes (Triantafyllou *et al.*, 2005). Considerable progress has been made recently in using 7 Tesla fMRI to study brain function at fine spatial resolution (Van der Zwaag *et al.*, 2013; Puckett *et al.*, 2017; Benson *et al.*, 2018; Huber *et al.*, 2020), however these protocols have been designed and optimised at a single research site and do not translate to studies where multiple sites are required to recruit the target population.

There has been a large recent expansion of 7 Tesla MRI infrastructure, driven by the release of the first commercially available clinical (CE-marked) 7 Tesla MRI system (U. S. Food & Drug Administration, 2017), opening up the possibility of wide translation of 7 Tesla to clinical applications. Further, recent work from the UK 7T Network (Clarke *et al.*, 2020; Rua *et al.*, 2020), German Ultra-high Field Imaging Network (Voelker *et al.*, 2016) and European Ultrahigh-Field Imaging Network in Neurodegenerative Diseases (Düzel *et al.*, 2019) have started to establish protocols for harmonising 7 Tesla MRI measurements across sites and scanner models with good reproducibility, providing the framework to disseminate techniques across sites.

In order to assess fMRI reproducibility across 7T sites an experimental design that exploits the advantages of ultra-high field is desirable, such as high spatial resolution. The study reported here investigates reproducibility of hand-digit representation measurements using 7 T fMRI across five centres, with measurements from three different 7 Tesla whole-body MRI systems from two different vendors. As part of the UK 7T Network’s “Travelling-Heads” study, 10 participants were scanned at 5 sites, including a hand-digit localization fMRI task (Sanchez-Panchuelo *et al.*, 2010; Kolasinski *et al.*, 2016). The dataset from a single subject has been released as part of the UK 7T harmonization paper (Clarke *et al.*, 2020) and this work has been presented in abstract format (Driver *et al.*, 2021; https://www.ismrm.org/21m/).

## Methods

Ten healthy subjects (32±6 years; 3 female; 1 left-handed) participated in the study. To assess inter-site repeatability, the same protocol was repeated at five sites, using three different 7 Tesla whole-body MRI systems, as detailed in *Table 1*. The same model of volume-transmit, 32-channel receive head coil (Nova Medical) was used at each site. Subjects were split into five pairs, with each pair returning to one site an additional four times (two subjects per site), to assess intra-site repeatability.

**Table 1:** MRI System specifications for each site

| # | Site | Vendor | Scanner Model | Gradient Performance |
| --- | --- | --- | --- | --- |
| 1 | Wolfson Brain Imaging Centre, University of Cambridge | Siemens | Magnetom Terra | 80 mT m<sup>-1</sup><br>200 mT m<sup>-1</sup> ms<sup>-1</sup> |
| 2 | Cardiff University Brain Research Imaging Centre, Cardiff University | Siemens | Magnetom 7T | 70 mT m<sup>-1</sup><br>200 mT m<sup>-1</sup> ms<sup>-1</sup> |
| 3 | Imaging Centre of Excellence, University of Glasgow | Siemens | Magnetom Terra | 80 mT m<sup>-1</sup><br>200 mT m<sup>-1</sup> ms<sup>-1</sup> |
| 4 | Sir Peter Mansfield Imaging Centre, University of Nottingham | Philips | Achieva 7T | 40 mT m<sup>-1</sup><br>200 mT m<sup>-1</sup> ms<sup>-1</sup> |
| 5 | Wellcome Centre for Integrative Neuroimaging (FMRIB), University of Oxford | Siemens | Magnetom 7T | 70 mT m<sup>-1</sup><br>200 mT m<sup>-1</sup> ms<sup>-1</sup> |

Two runs of a visually cued, travelling wave somatomotor task were performed with the dominant hand. This entailed a visually paced sequential 1 Hz button press of 8 s blocks of digit movement, cycling across digit blocks from index finger (D2) to little finger (D5) in a “forward” run and from D5 to D2 in a “reverse” run (32s per four digits), with 8 cycles per run (*Figure 1*).

**Figure 1:**
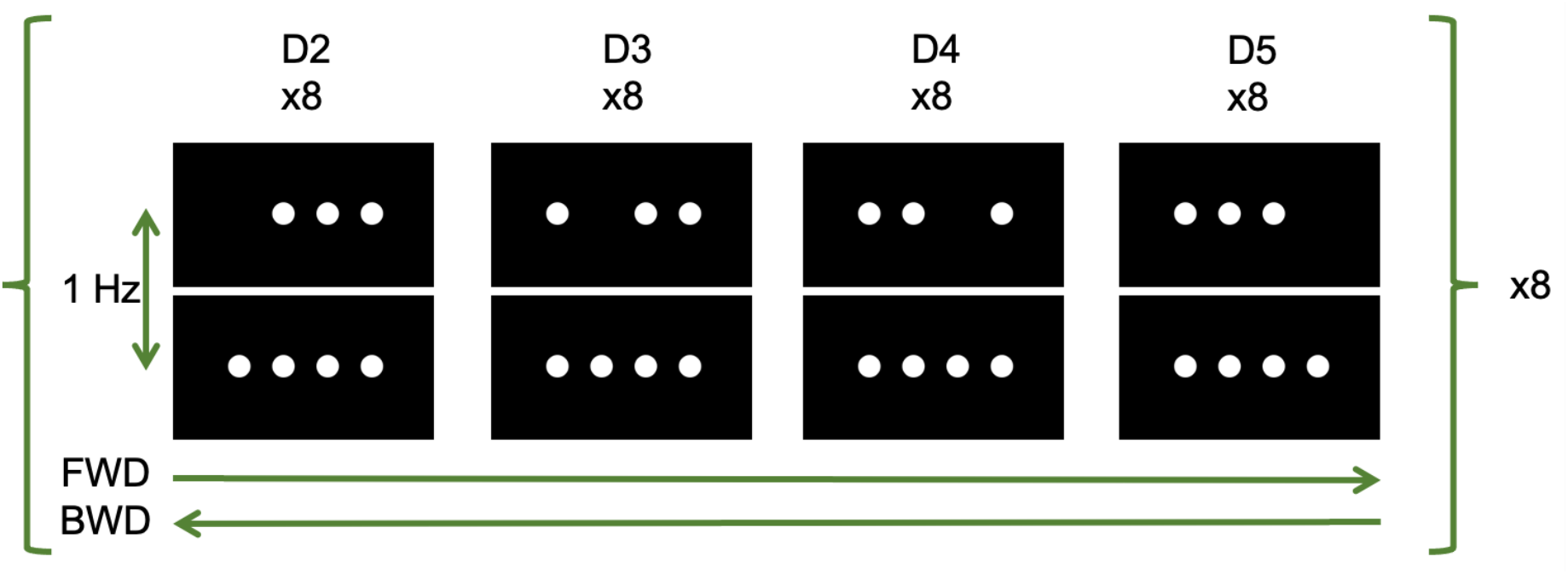
Hand digit localization task design. Participants were cued to press a button in time with a flashing circle, corresponding to each digit. FWD – “forward” run, from index finger (D2) to little finger (D4); BWD – “reverse” run, from little finger to index finger.

MRI calibration steps of radiofrequency transmit power calculation (using a 3DREAM B_0_^+^ map) and B_0_ shimming were matched across sites, using the protocol that we introduced in our previous work (Clarke *et al.*, 2020). Gradient-echo EPI protocol: 1.5mm isotropic resolution, TR=2s, TE=25ms, echo spacing 0.68/0.78ms Siemens/Philips, 34/28 slices Siemens/Philips. An in-plane acceleration factor of 2 was used, with GRAPPA for Siemens sites and SENSE for the Philips site. Spin-echo EPI scans were acquired with matched and reversed phase-encode direction for distortion correction (Driver *et al.*, 2018; Clarke *et al.*, 2020). A T_1_-weighted MPRAGE (0.7mm isotropic, TR/TE/TI=2200/3.05/1050ms, FA=7°) dataset was acquired for realignment and cortical flattening. A T2*-weighted 2D FLASH (0.75×0.75×1.5mm^3^, TE=10ms, TR=1100ms) dataset was acquired as an intermediate registration step between the EPI and T_1_-weighted MPRAGE.

Data was motion corrected using FSL MCFLIRT (Jenkinson *et al.*, 2002), distortion corrected with FSL TOPUP (Andersson, Skare and Ashburner, 2003), and temporal filtered (100s cut-off) using FSL FEAT (Smith *et al.*, 2004). A Fourier-based travelling wave analysis was performed using mrTools (Gardner *et al.*, 2018) to calculate voxel-wise phase and coherence of the BOLD response. Image registration was performed using mrTools (Gardner *et al.*, 2018). Whole-hand activation regions (FDR-corrected p < 0.05) were compared across sessions using Dice’s overlap coefficient (Dice). An intersection mask was formed for each subject by including voxels identified as active across all five sites. Individual digit maps were formed within this intersection mask by dividing the phase maps into four equal π/2 portions. Digit overlap across sessions was compared using the Dice similarity coefficient. Intra- and inter-site reproducibility were compared using Bonferroni-corrected paired t-tests applied to the mean Dice coefficient across repeated sessions in the same site and across sites, respectively.

## Results

Temporal SNR was similar across the five sites, with tSNR = 45±4/43±4/46±5/45±4/44±3 (mean±standard deviation across subjects) for sites 1-5, respectively and 1-way ANOVA F(4,45)=0.89, p = 0.48, testing for differences in tSNR across sites. The comparison of tSNR was only considered across the 5 inter-site sessions, since including the repeat intra-site sessions would skew results to individual subjects, with only 2 subjects per site participating in the repeat sessions.

*Figure 2* presents example conjunction maps for five subjects, showing the number of sessions with overlapping whole-hand activation regions, across repeats at the same site (intra-site), or across sites (inter-site). These datasets were chosen to present intra-site data from each of the five sites.

**Figure 2:**
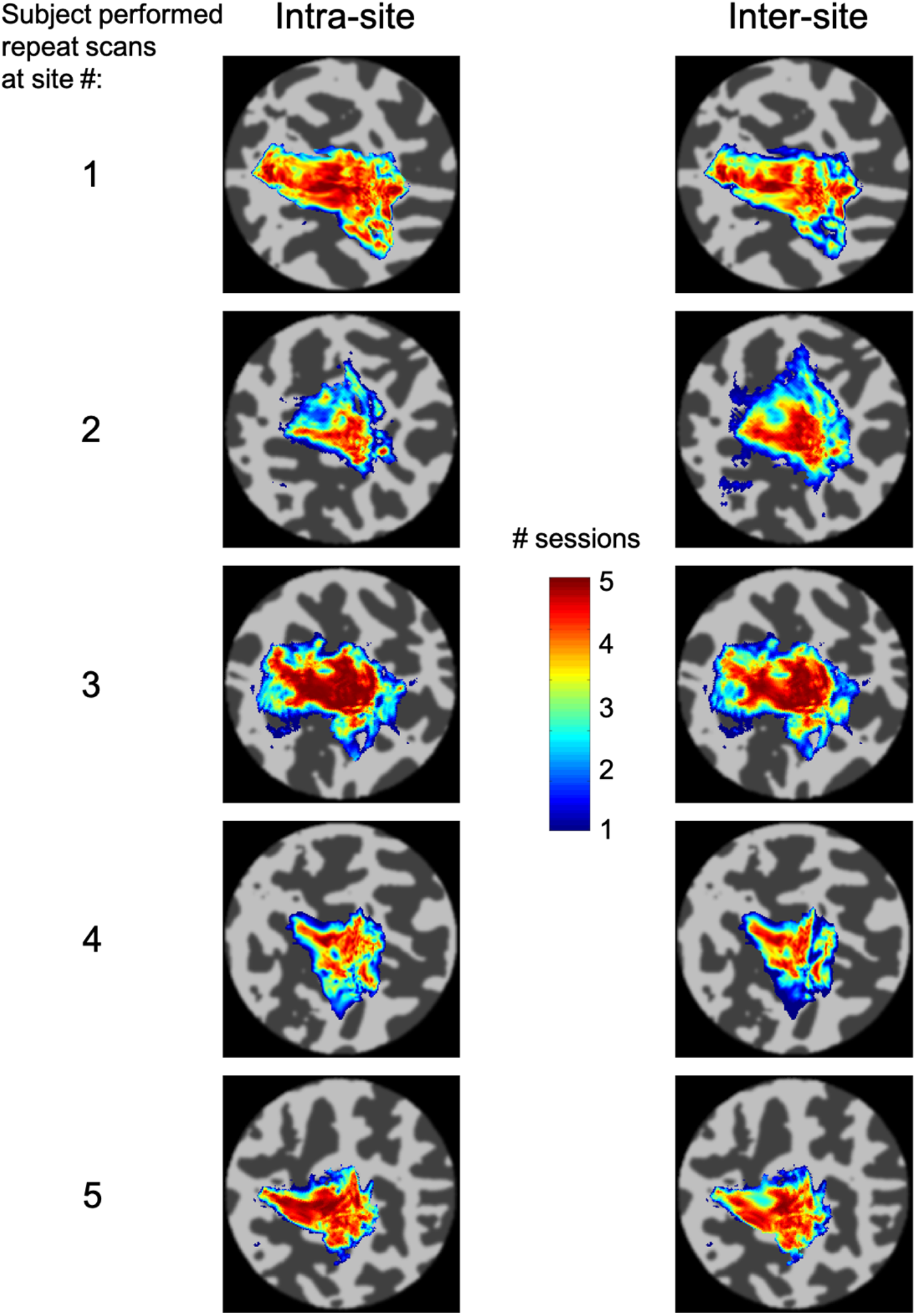
Conjunction maps for five subjects showing the overlap of the hand region between sessions for both repeat sessions for a single site (intra-site) or across sites (inter-site). The five subjects were chosen to represent intra-site measures for each site.

Phase (digit) maps are shown in *Figure 3*, with all sessions from a single subject shown (*Fig. 3a*) and all ten subjects’ phase maps for a single site (*Fig. 3b*). The spatial distribution of the digit representations are consistent across sites for a single subject, whereas there is a high degree of inter-subject variability. *Figure 4* shows both intra-site and inter-site conjunction maps for each digit, with a high degree of similarity between intra-site and inter-site for individual digits.

**Figure 3:**
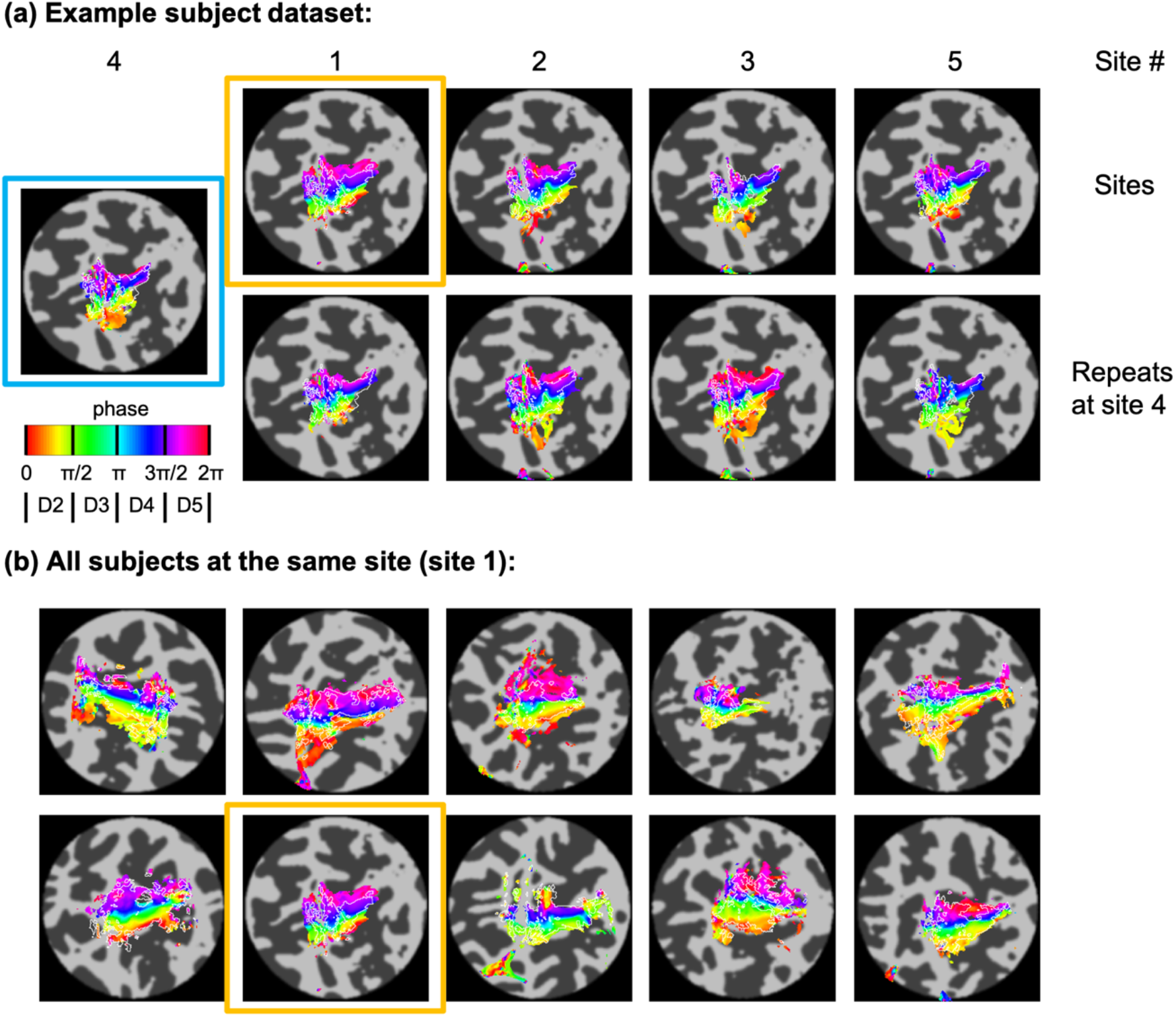
Phase maps and associated digit maps (D2-D5) defined by dividing phase into π/2 portions. Overlay shown for p_FDR_ < 0.05. Intersection mask outline shown in white (p_FDR_ < 0.05 hand region for all five sites). (a) Phase maps from all sessions for a single subject; (top row) all sites; (bottom row) repeat sessions at site 4. Blue border indicates data in both intra-site and inter-site analysis. (b) Phase maps from all 10 subjects for site 1. Orange border matches the session in (a).

**Figure 4:**
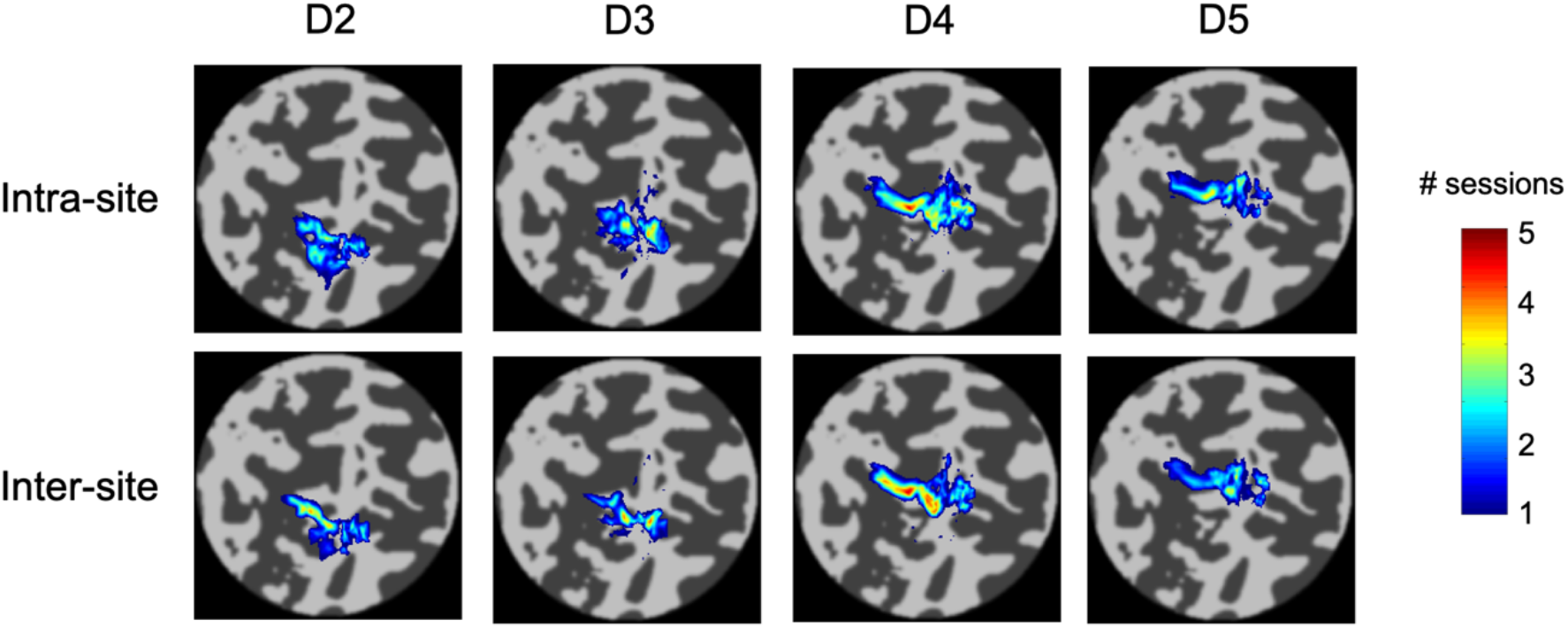
Conjunction maps for each digit (index finger - D2; little finger – D5) for a single subject, showing the overlap of the digit region between sessions for both repeat sessions for a single site (intra-site) or across sites (inter-site). The subject performed repeated sessions at site 4.

Dice coefficients for the whole hand region are higher for intra-session measures from the individual sites compared to inter-site measures (*Figure 5*; t(9) = −5, p_corr_ = 0.002). There is also a similar trend in the individual digit maps for intra-site Dice coefficients to be greater than inter-site (*Fig. 5*), but this only survives Bonferroni correction (p_corr_<0.05) in D3 and D4.

**Figure 5:**
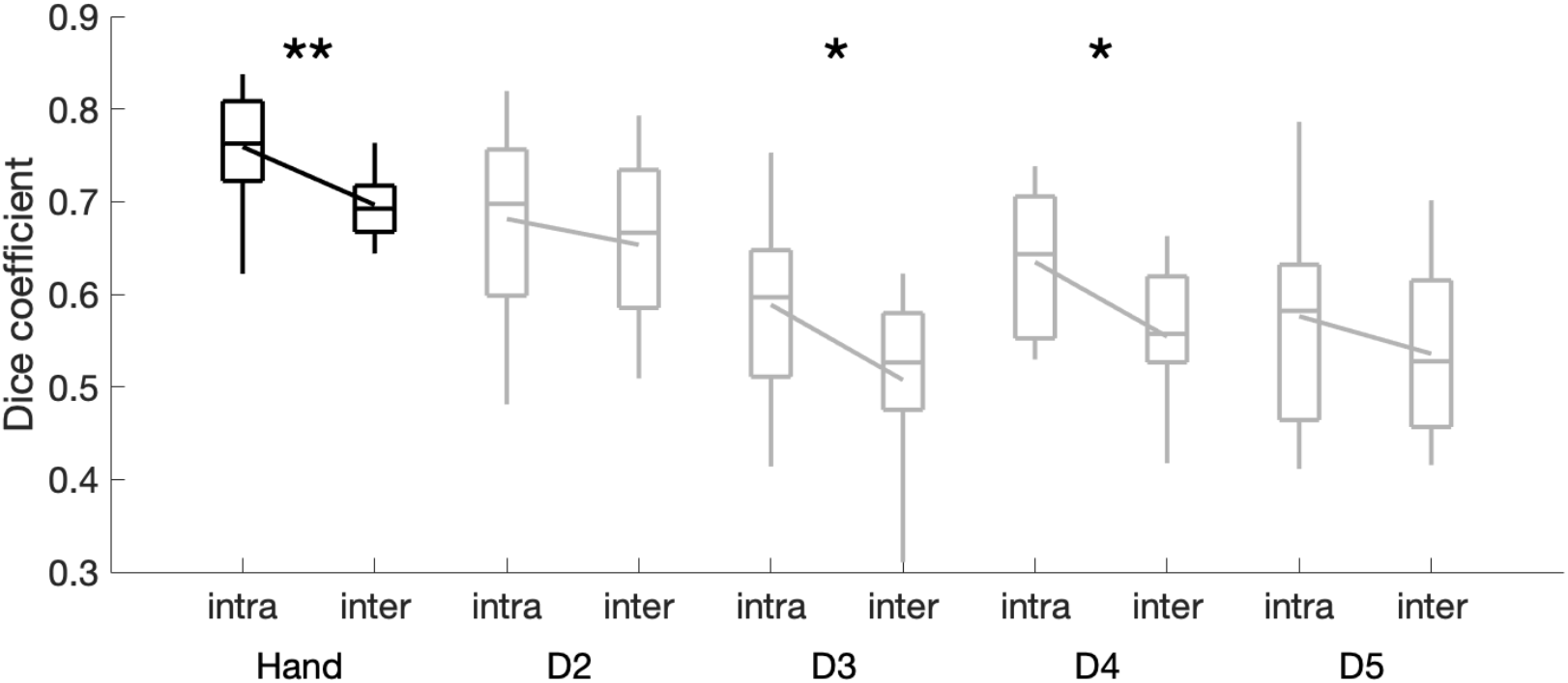
Boxplots comparing intra- and inter-site Dice coefficients for the whole hand region and each digit. Lines connecting intra/inter pairs compare their respective means. **p<0.005; *p<0.05 (Bonferroni-corrected)

## Discussion

This study demonstrates good reproducibility of fMRI digit maps across five, 7 T sites, using three different models of whole-body MRI system from two manufacturers. We did not harmonize the image reconstruction, coil combination, or parallel acquisition methods. Simple protocol harmonization, such as matching EPI echo train duration, in-plane acceleration factor, acquisition matrix, TE and TR, is sufficient to achieve good reproducibility across MRI systems, without needing to match gradient waveforms and radiofrequency pulse shapes.

The 1.5 mm isotropic spatial resolution fMRI data acquired here was able to resolve inter-subject differences in hand digit representations, consistent with previous observations (Kolasinski *et al.*, 2016). By repeating measurements across sites in the same participants, we were able to isolate inter-site variability from inter-subject differences. Qualitatively, inter-site and intra-site variability is small compared to these inter-subject differences. We show that the inter-site Dice overlap was smaller than intra-site. However, this difference in Dice coefficients corresponds to ~5% difference in overlap in the regions between intra- and inter-site comparisons, which is small compared to the inter-session variability. Inter-site Dice values are similar to those reported across sessions in an intra-site study (Kolasinski *et al.*, 2016). These findings can be used to power sample sizes required for future multi-site fMRI studies involving hand digit localization and show that such studies can gain from a larger participant pool available across multiple sites, with the increase in measurement variability from multisite measurements being small compared to the test-retest variability within a single site.

The region of interest definition was based on the coherence of the data studied, including voxels which were defined as active at all five sites. This approach risks missing novel features from individual sessions. Future work will define regions of interest based on a probabilistic atlas of hand digit areas (O’Neill *et al.*, 2020). This approach will yield a broader region, including areas that may not be active in individual sessions, leading to increased variability in digit maps, however it will provide a region of interest definition that is independent of the data studied.

## Conclusion

High resolution fMRI studies can be performed across multiple sites, with inter-site factors contributing. The inter-site variability measured here, in the form of Dice coefficients, can be used to inform future study designs and sample size calculations for multi-site somatomotor mapping studies.

## Acknowledgements and Funding

The UK7T Network and this work was funded by the UK’s Medical Research Council. [MR/N008537/1].

## Centre funding

The Wellcome Centre for Integrative Neuroimaging is supported by core funding from the Wellcome Trust (203139/Z/16/Z).

Cardiff University Brain Research Imaging Centre is supported by the UK Medical Research Council (MR/M008932/1) and the Wellcome Trust (WT104943).

This research was supported by the NIHR Cambridge Biomedical Research Centre (BRC-1215-20014). The views expressed are those of the author(s) and not necessarily those of the NIHR or the Department of Health and Social Care. The Cambridge 7T MRI facility is cofunded by the University of Cambridge and the Medical Research Council (MR/M008983/1).

## Individual funding

CTR is funded by a Sir Henry Dale Fellowship from the Wellcome Trust and the Royal Society [098436/Z/12/B].

